# DOA/CLK2 phosphorylates Fl(2)d/WTAP to enhance m^6^A mRNA methyltransferase complex activity

**DOI:** 10.1101/2024.11.25.625202

**Authors:** David W. J. McQuarrie, Wei Bian, Matthias Soller

## Abstract

*N*^6^-methyladenosine (m^6^A) is the most abundant internal modification in messenger RNA (mRNA) providing an essential layer to the control of gene expression important to many biological processes. A megadalton writer complex deposits m^6^A methyl marks that are then read by YTH domain containing proteins. Deposition of m^6^A is thought to be regulated by cellular signalling but how remains uncertain. We identified the kinase DOA, a highly conserved homologue of human CLK2 by using the essential role of m^6^A in the *Drosophila* Sxl auto-regulation for sex determination and dosage compensation. We show that DOA kinase is required for m^6^A deposition and *Sxl* alternative splicing. Overexpression of DOA can compensate writer complex insufficiency and rescue lethality of m^6^A in the context of *Sxl* mis-splicing, suggesting a key role in regulating activity of the m^6^A complex. Through genetic interaction experiments, we identify a phospho-site at the end of a conserved helical structure in Fl(2)d as DOA target. CLK2 phosphorylates the same helix in WTAP, the human homologue of Fl(2)d. CLK2 expands regulatory capacity through additional phospho-sites and changing of the *Drosophila* site to a phosphomimetic-like glutamine, overall important for m^6^A directed regulation of dosage compensation genes in human cells.

## Background

Following transcription, various nucleotides of mRNAs can be modified to add an essential layer to the regulation of gene expression (Roignant and Soller 2017; Anreiter et al. 2020; Anreiter et al. 2022). Among mRNA modifications, *N*^6^-methyladenosine (m^6^A) is the most abundant internal modification in eukaryotes (Balacco and Soller 2019; Zaccara et al. 2019). The m^6^A modification plays a pivotal role in regulating mRNA metabolism and is central to numerous biological processes, including cell differentiation, DNA damage response circadian rhythm, neurogenesis and sex determination (Fustin et al. 2013; Lence et al. 2016; Lence et al. 2017; Roignant and Soller 2017; Elvira-Blázquez et al. 2024). Dysregulation of m^6^A has been implicated in numerous diseases, metabolic disorders, neurodegeneration and various forms of cancer (Fitzsimmons and Batista 2019; Sun et al. 2019; Han et al. 2020; Jiang et al. 2020; Qin et al. 2020).

The m^6^A epimark is deposited by a dedicated and evolutionarily highly conserved megadalton methyltransferase “writer” complex, consisting of two sub-complexes: the m^6^A– METTL complex (MAC) made up of a methyltransferase-like 3 (METTL3) and METTL14 heterodimer (Balacco and Soller 2019; Zaccara et al. 2019; Ensinck et al. 2023). In MAC, METTL3 is the catalytic subunit and interacts with the catalytically dead METTL14, but MAC without the auxiliary proteins has little activity (Bokar et al. 1997; Liu et al. 2014). The m^6^A-METTL-associated complex (MACOM) contains the auxiliary subunits Wilms tumour 1- associated protein (WTAP), Vir-like m^6^A methyltransferase-associated (KIAA1429/VIRMA), HAKAI/CBLL1, RNA-binding motif 15 (RBM15) and its paralog RBM15B, and Fl(2)d-associated complex component (FLACC/ZC3H13) (Balacco and Soller 2019; Zaccara et al. 2019). These are conserved in *Drosophila* as Mettl3, Mettl14, Female-lethal (2)d (Fl(2)d/WTAP), Virilizer (Vir/VIRMA), Hakai, Spenito (Nito/RBM15), and Flacc (Lence et al. 2017; Knuckles et al. 2018; Balacco and Soller 2019; Bawankar et al. 2021; Wang et al. 2021). Cryo-electron microscopy (cryo-EM) revealed a partial MACOM complex and the interaction interface of METTL3 and METL14 in the MAC heterodimer has been determined by X-ray crystallographic structural studies (Wang et al. 2016; Su et al. 2022). In addition, the complex array of interactions in the core-MACOM consisting of Fl(2)d, Vir, Flacc and Hakai are required for its stability and m^6^A methyltransferase activity (Bawankar et al. 2021; Wang et al. 2021).

A further level of regulation is imposed on m^6^A deposition by post-translational modifications. The Mitogen-Activated Protein Kinase (MAPK) pathway has been shown to phosphorylate METTL3 and WTAP through ERK (Sun et al. 2020). Ubiquitin Specific Peptidase 5 (USP5) can deubiquitinate METTL3 and WTAP (Sun et al. 2020). The stepwise combination of these post-translational modifications stabilises the m^6^A methyltransferase complex (Sun et al. 2020). In addition, WTAP phosphorylation has been shown to lead to its aggregation, while in contrast, dephosphorylation by PPP4 enhances phase separation (Cai et al. 2024). The MACOM member Hakai is an E3 ubiquitin ligase, however, global ubiquitination levels are unaffected by Hakai knockdown, suggesting a more specific role in the regulation of methyltransferase complex activity (Bawankar et al. 2021).

Functions of the m^6^A methyltransferase complex have been associated with various sexual phenotypes including X-chromosome inactivation in humans, sex determination and dosage compensation in *Drosophila* and meiosis in yeast (Roignant and Soller 2017). In *Drosophila*, the master regulator of sex determination Sex-lethal (Sxl) is only expressed in females. Sxl autoregulates through alternative splicing by supressing inclusion of a male-specific exon with a translational stop codon. In this regulated intron, Sxl binds to sites flanking the splice sites of the male exon. In the absence of m^6^A, or its nuclear reader YTHDC1, suppression of the male exon is insufficient leading to reduced levels of Sxl, which then impacts on Sxl’s second role in dosage compensation through translational suppression of *msl-2* expression in females. MSL-2 is only present in males and through the MSL complex doubles gene-expression of the single male X chromosome. Now, if m^6^A is missing, Sxl levels are reduced leading to insufficient dosage compensation, which compromises female fitness often resulting in lethality. As indicated from saturating genetic screens (Schutt and Nothiger 2000), the m6A writer complex and its reader have a particularly critical role in *Sxl* alternative splicing allowing genetic identification of additional regulators of this pathway (Haussmann et al. 2016; Lence et al. 2016; Knuckles et al. 2018; Bawankar et al. 2021; Wang et al. 2021).

Sexual transformations have been observed in mutants of the kinase Darkener of Apricot (DOA) (Du et al. 1998). These phenotypes have been associated with phosphorylation of the key sex determination protein Transformer (TRA), a downstream target of Sxl (Du et al. 1998; Yun et al. 2000; Kpebe and Rabinow 2008a; Kpebe and Rabinow 2008b; Zhao et al. 2015). TRA is an RNA-binding protein of the SR protein family and DOA phosphorylates the SR domain to impact of alternative splicing of the sex determination gene *doublesex* into male- and female-specific transcript factor isoforms (Du et al. 1998). DOA is the highly conserved homologue of human CDC-like kinase 2 (CLK2, Fig. 1A, Supplementary Fig. 1), which has been shown to regulate alternative splicing in humans through phosphorylation of SR proteins (Hu et al. 2024; Liu et al. 2024).

**Fig. 1:**
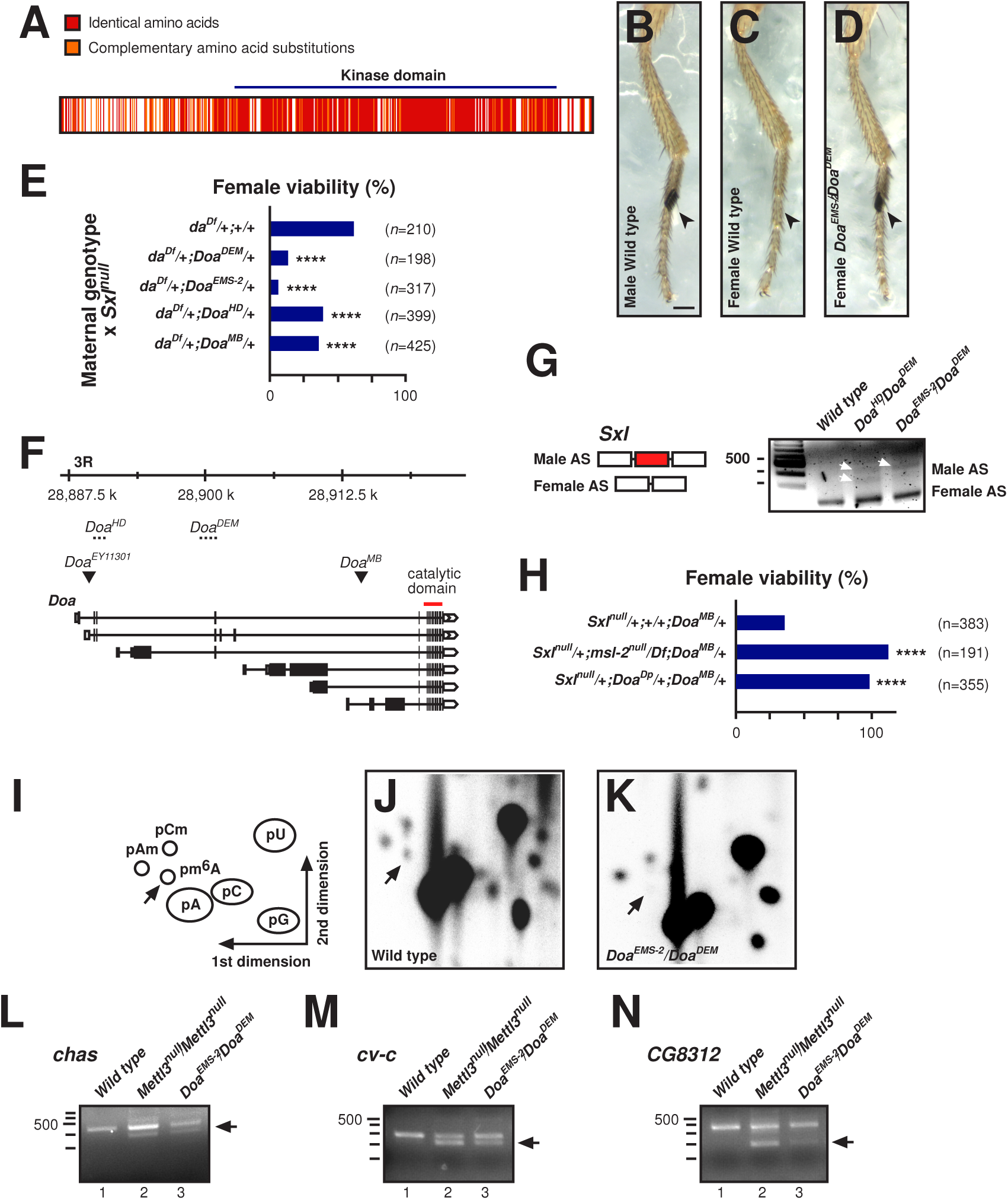
DOA is required for female dosage compensation and m^6^A methylation. A) Amino acid alignment of DOA and CLK2 between *D. melanogaster* and humans, respectively. Identical amino acids are highlighted (red), complementary amino acid substitutions (orange), and non-complementary amino acid substitutions (unmarked) are indicated. B-D) Front legs of indicated genotypes. The scale bar (D) is 100 µm. E) Quantification of female viability for the indicated genotypes when crossed to *Sxl*^*7B0*^ males.
Heterozygous *Doa* alleles significantly reduce female viability in a *Sxl* and *da* heterozygous null background. Statistically significant differences calculated from chi-squared tests following FDR correction are indicated by asterisks (**** p≤0.0001). F) *Doa* gene model with alleles used in this study. Lethal *Doa*^*HD*^ and *Doa*^*DEM*^ are linked to inversions and *Doa*^*EMS-2*^ has not been mapped molecularly. *Doa*^*MB*^and *Do*^*BEY11301*^ are viable transposon inserts. The catalytic domain is indicated as a red line. G) RT-PCR analysis of male isoform *Sxl* splicing in females between exons 2 and 4 for wild- type, *Doa*^*HD*^*/Doa*^*DEM*^, and *Doa*^*EMS-2*^*/Doa*^*DEM*^ head/thorax. H) Quantification of female viability for the indicated genotypes when crossed to *Sxl*^*7B0*^ males. Complete loss of *msl-2* and duplication of *Doa* rescued female viability in *Doa* mutants in a *Sxl* null background. Statistically significant differences calculated from chi-squared tests following FDR correction are indicated by asterisks (**** p≤0.0001). I-K) Schematic representation of a TLC results depicting non-methylated and methylated nucleotide positions (I), and TLCs displaying m^6^A in wild-type (J) and *Doa*^*EMS-2*^*/Doa*^*DEM*^female adults (K). L-M) RT-PCR of alternative splicing for *chas* (L), *cv-c* (M), *and CG8312* (N) in wild-type,
*Mettl3*^*null*^, and *Doa*^*EMS-2*^/*Doa*^*DEM*^ female adults.

Here, we identify a novel regulatory role for DOA/CLK2 through its role in female-specific *Sxl* splicing in regulating m^6^A complex activity. *Doa* genetically interacts with *Sxl* as found typical for m^6^A writer complex components, e.g. lethality can be rescued by preventing ectopic dosage compensation in females. Moreover, overexpression of DOA rescues lethality of m^6^A writer complex insufficiency in the context of *Sxl* mis-splicing, suggesting a key role in regulating activity of the m^6^A complex. Consistent with such key role, *Doa* mutants are devoid m^6^A in mRNA. Instructed by genetic interaction experiments, we identify Fl(2)d as the phosphorylation target, where thyronine 301 (T301) at the end of the conserved helical structure is phosphorylated by DOA. Analysis of CLK2 function in human cells reveals a conserved role in m^6^A methylation by phosphorylation of WTAP. Here, CLK2 adopts a new regulatory capacity through additional phospho-sites and fixation of the phosphomimetic-like amino acid glutamine in WTAP at the *Drosophila* DOA-regulated site.

## Results

### DOA regulates female *Sxl* splicing for robust dosage compensation and female differentiation

To identify regulators of the m^6^A complex, we focused on genes whose loss of function results in sexual transformation phenotypes. In one such gene, *Doa*, we observed 10% of *Doa^EMS-2^*/*Doa^DEM^* females (n=30) displaying ectopic sex combes (Fig. 1B-D) (Du et al. 1998; Yun et al. 2000; Kpebe and Rabinow 2008b; Hoedjes et al. 2022). To test whether *Doa* is required for *Sxl* autoregulation, we used a genetically sensitized background containing one copy of *Sxl* and *daughterless* (*da*), which is involved in *Sxl* transcription, to reduce *Sxl* levels (Bawankar et al. 2021). In the progeny of a cross between *da^Df^*/+; *Doa*/+ females and *Sxl^7B0^* null males, most females died for the strongest *Doa^DEM^* and *Doa^EMS-2^* alleles, while in the weaker alleles *Doa^HD^* and *Doa^MB^* female viability was less affected. (Fig. 1E and F). In addition, we also observed male specific abdominal pigmentation in *Doa* sensitised females, indicative of sex determination defects, which is cell-autonomous (Supplementary Fig. 2).

Next, we analysed *Sxl* alternative splicing in *Doa* heteroallelic mutants (*Doa^EMS-2^*/*Doa^DEM^* and *Doa^HD^*/*Doa^DEM^* females). In *Doa* mutant females, inclusion of the male-specific exon was increased (Fig. 1G).

To validate that female lethality in *Sxl* sensitized *Doa* mutants is due to mis-regulation of dosage compensation, we removed *msl-2* in these females. In this condition, *Doa^MB^*female viability was restored (Fig. 1H). Lethality was also rescued by providing a duplication of the *Doa* locus (*Doa^Dp^*, Fig 1G). Taken together, these results identify DOA as a genuine regulator of *Sxl* through its role in sex determination and dosage compensation.

### DOA is required for m^6^A methylation and m^6^A dependent splicing events

In addition to *Sxl*, m^6^A mRNA methylation and decoding by YTHDC1 is required for robust sex determination and dosage compensation through its role in supporting Sxl in autoregulation. To test whether DOA acts through the m^6^A pathway we determined m^6^A levels in poly(A) RNA of *Doa^EMS-2^*/*Doa^DEM^*mutants via 2D thin layer chromatography (TLC). We observed that m^6^A essentially is absent in *Doa* mutants (Fig. 1I-K).

To further substantiate DOA’s role in the m^6^A pathway, we analysed alternative splicing of *chas*, *cv-c*, and *CG8312* which are dependent on Mettl3 deposition of m^6^A (Haussmann et al. 2016). Here we observed splicing changes in all genes for *Mettl3* and *Doa* mutants compared to the control (Fig. 1L-N).

Expression of m^6^A writers and readers is increased in the brain and their loss results in neurological defects (Haussmann et al. 2016; Lence et al. 2016; Kan et al. 2017; Worpenberg et al. 2021). At neuro-muscular junctions in third instar larvae, *Mettl3^null^* mutants have more synapses, which we also observed in *Doa^EMS-2^*/*Doa^DEM^* mutants compared to the control (Supplementary Fig. 3). Taken together these results indicate that DOA regulates m^6^A methyltransferase complex activity.

### DOA regulates m^6^A methyltransferase complex activity and drives its tissue-specific regulation

We previously observed that Hakai affects *Sxl* splicing differentially in head/thorax compared to abdomen when sensitised by loss of *vir* indicating regulation of the m^6^A pathway by cellular signalling (Bawankar et al. 2021). To identify how DOA contributes to tissue-specific regulation of the m^6^A pathway, we removed one copy of *Doa* in a *vir^2F^*/*vir^ts^* genetically sensitised background. This condition resulted in male pigmentation in ∼50% of *Doa^MB^* females in the sensitised homozygous *vir^2F^*/ *vir^ts^*background (Fig. 2A-F, Supplementary Fig. 4, Supplementary results). Further, we noted some females with male sex combs and an intersex appearance in the *vir^2F^*/ *vir^ts^* background with one copy of *Doa* removed (Fig. 2B). Likewise, when we analysed *Sxl* splicing in these *vir^2F^*/ *vir^ts^* ;*Doa/+* females (Fig. 2C), we found tissue-specific changes between the head/thorax and abdomen, where the male *Sxl* splice isoform became dominant in addition to intermediary spliced products (Fig. 2F).

**Fig. 2:**
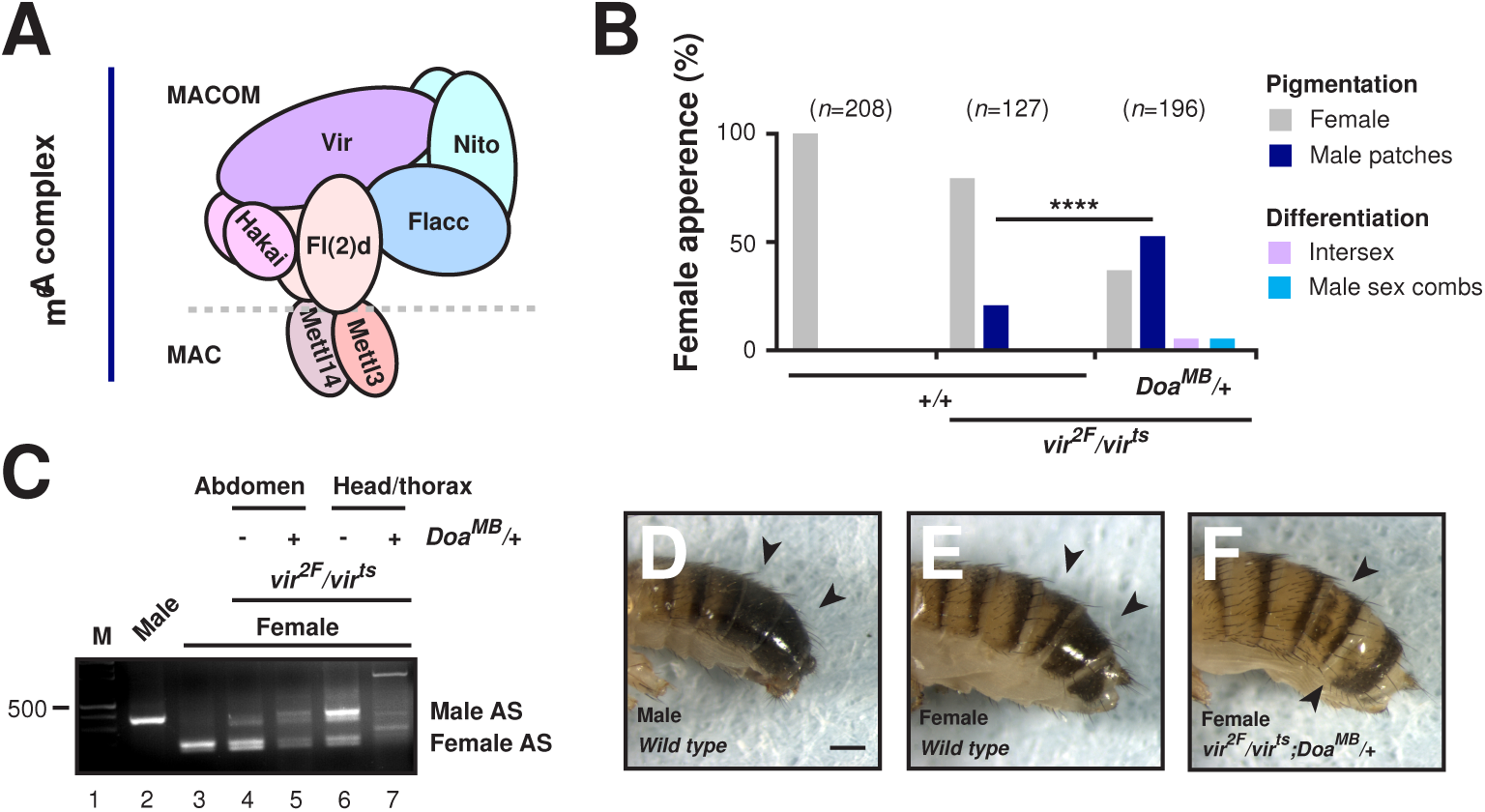
DOA regulates tissue-specific m^6^A complex activity. A) Schematic representation of the m^6^A methyltransferase writer complex. MAC and MACOM subcomplexes are indicated and divided by the dashed line. B) Quantification of male-specific pigmentation and sexual development in wild-type, *vir*^*2F*^/ *vir*^*ts*^, and *vir*^*2F*^/ *vir*^*ts*^; *Doa*^*MB*^*/*+ females. Females with a *vir*^*2F*^/ *vir*^*ts*^ background showed low levels of male pigmentation which was increased in *vir*^*2F*^/ *vir*^*ts*^ females heterozygous for *Doa*^*MB*^. C) RT-PCR analysis of male isoform *Sxl* splicing between exons 2 and 4 for male and female wild-type flies, and *vir*^*2F*^/ *vir*^*ts*^, and *vir*^*2F*^/ *vir*^*ts*^; *Doa*^*MB*^*/*+ female abdomen and head/thorax. D-F) Representative images of wild-type male and female pigmentation (D and E) and *vir*^*2F*^/ *vir*^*ts*^; *Doa*^*MB*^*/*+ male-specific pigmentation in females (F). The scale bar (D) is 200 µm.

To further substantiate DOA regulation of m^6^A writer complex activity, we wanted to test whether overexpression of DOA could increase m^6^A complex activity. For this experiment we made use of low female viability when one copy of *Mettl3* is removed in a sensitised *da* and *Sxl* background. When we overexpressed DOA in this genetic background of low Mettl3 levels, female viability is restored, which is explained by increased activity of the Mettl3 complex (Fig. 3A). When both copies of Mettl3 are removed no rescue is observed which is expected if DOA acts through Mettl3. We further confirmed this finding by removal of one copy of the other m^6^A complex members *Flacc*, *nito*, and *vir*, where DOA overexpression also restored female viability (Fig. 3A). For the *fl(2)d UASDoa* combination, crosses did not propagate indicating genetic interaction.

**Fig. 3:**
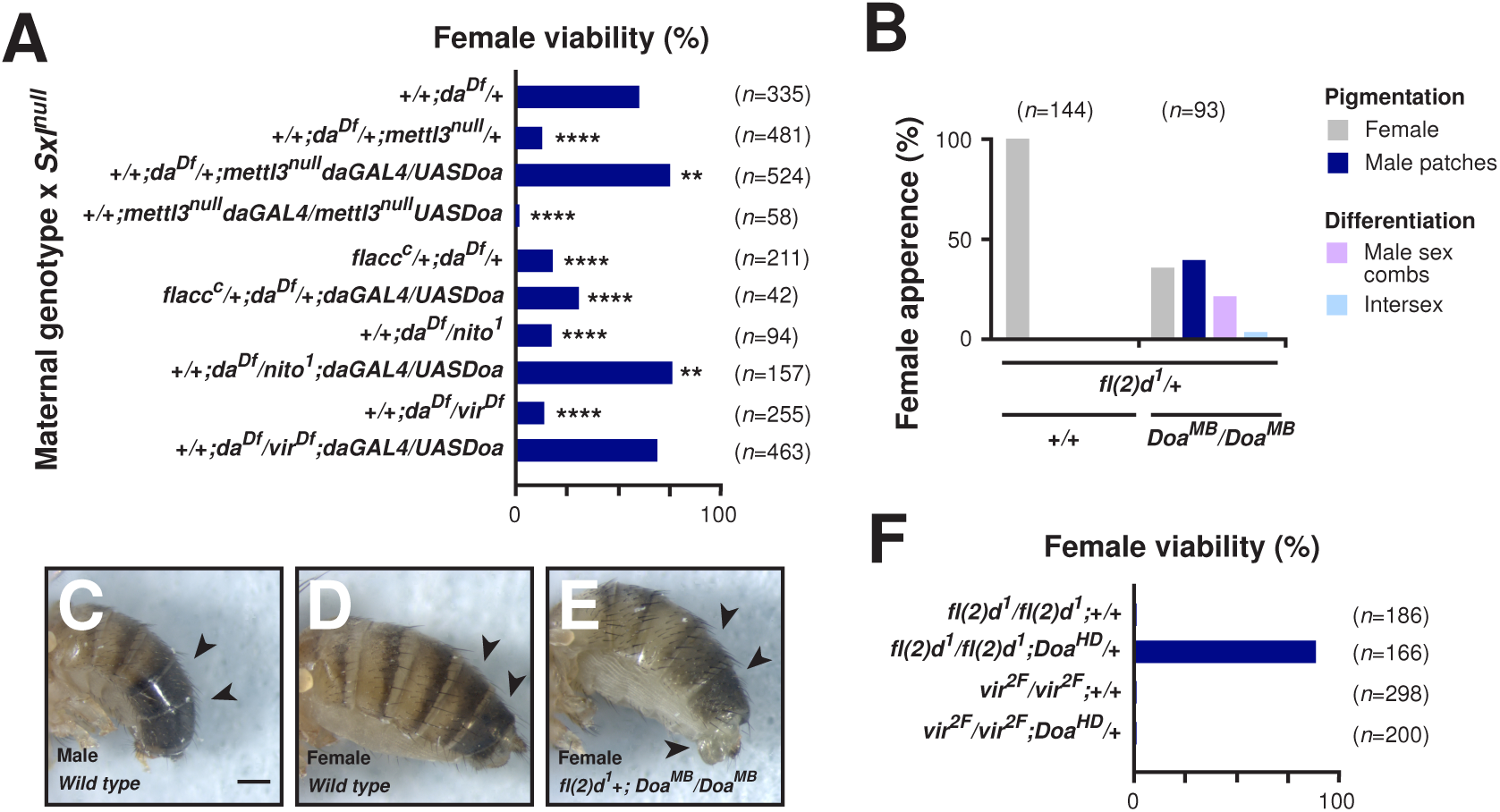
DOA overexpression rescues female lethality caused by low m^6^A writer concentrations through Fl(2)d. A) Quantification of female viability for the indicated genotypes when crossed to *Sxl*^*7B0*^ males. Overexpression of *Doa* rescues female viability for loss of one copy of an m^6^A methyltransferase writer in a *Sxl* and *da* heterozygous null background, but not in a homozygous *Mettl3*^*null*^ background. Statistically significant differences calculated from chi- squared tests following FDR correction are indicated by asterisks (** p≤0.01, **** p≤0.0001). B) Quantification of male-specific pigmentation and sexual development in *fl(2)d*^*1*^/ +, *fl(2)d*^*1*^/+; *Doa*^*MB*^*/Doa*^*MB*^, and *fl(2)d*^*1*^/*Doa*^*Dp*^; *Doa*^*MB*^*/Doa*^*MB*^ females. Females with a *fl(2)d*^*1*^/+; *Doa*^*MB*^*/Doa*^*MB*^ background displayed male-specific pigmentation, male sex combs, and intersex phenotypes. C-E) Representative images of wild-type male and female pigmentation (C and D) and *fl(2)d*^*1*^/+; *Doa*^*MB*^*/Doa*^*MB*^male-specific pigmentation and differentiation in females (E). The scale bar (C) is 200 µm. F) Rescue of female lethality of the female-lethal *fl(2)d*^*1*^ allele, but not the *vir*^*2F*^ allele through removal of one copy of *Doa*.

Removal of the core MACOM complex members Fl(2)d, Vir, Flacc, or Hakai, but not Nito, leads to destabilisation of the m^6^A writer complex, evident by reduced levels of these members (Bawankar et al. 2021; Wang et al. 2021). To analyse whether DOA is required to stabilise the m^6^A writer complex we generated mitotic clones for the lethal *Doa^EMS-2^* allele in wing discs and stained for Fl(2)d. Here, we did not see a discernible reduction of Fl(2)d levels (Supplementary Fig. 5).

These findings further argue that DOA genuinely regulates m^6^A writer complex activity and strengthens the view that m^6^A deposition is regulated by cellular signalling rather than being a passive process (Uzonyi et al. 2023).

### DOA regulates m^6^A methyltransferase complex activity through Fl(2)d

To genetically identify the target of DOA in the m^6^A methyltransferase complex, we used the homozygous viable *Doa^MB^* allele as a genetically sensitised *Doa* background. We then removed one copy of each of the m^6^A writer complex components (*Mettl3*, *Mettl14*, *fl(2)d*, *Hakai*, *vir*, *nito*, *Flacc*) in the absence of *Doa* to see whether this condition induces sexual transformations. However, we only observed sexual transformations for removal of *fl(2)d* resulting in the majority of females displaying male pigmentation, male sex combs and/or intersex characteristics (Fig. 3B-E).

Female-lethal alleles have been identified in genetic screens for *fl(2)d* and *vir*. Female lethality can be rescued by removal of m^6^A writer Mettl3 (Haussmann et al. 2016; Knuckles et al. 2018; Bawankar et al. 2021). When we removed one copy of *Doa*, female viability of *fl(2)d^1^*, but not *vir^2F^*was rescued (Fig. 3F) indicating that Fl(2)d is the target of DOA.

### DOA/CLK2 phosphorylates the C-terminus of the extended alpha-helix in Fl(2)d/WTAP

To identify the DOA phosphorylation sites in Fl(2)d, we expressed N- and C-terminal fragments of Fl(2)d in *E. coli* to produce recombinant proteins (Fig. 4A). We then incubated these fragments with recombinant human CLK2 and P^32^-gamma-ATP we observed that the C-terminal region was phosphorylated (Fig. 4A). Mass spectrometry analysis of post-translational modifications on the Fl(2)d C-terminal fragment revealed phosphorylation of threonine 301 (T301), which is conserved in insects (Fig. 4B and C).

**Fig. 4:**
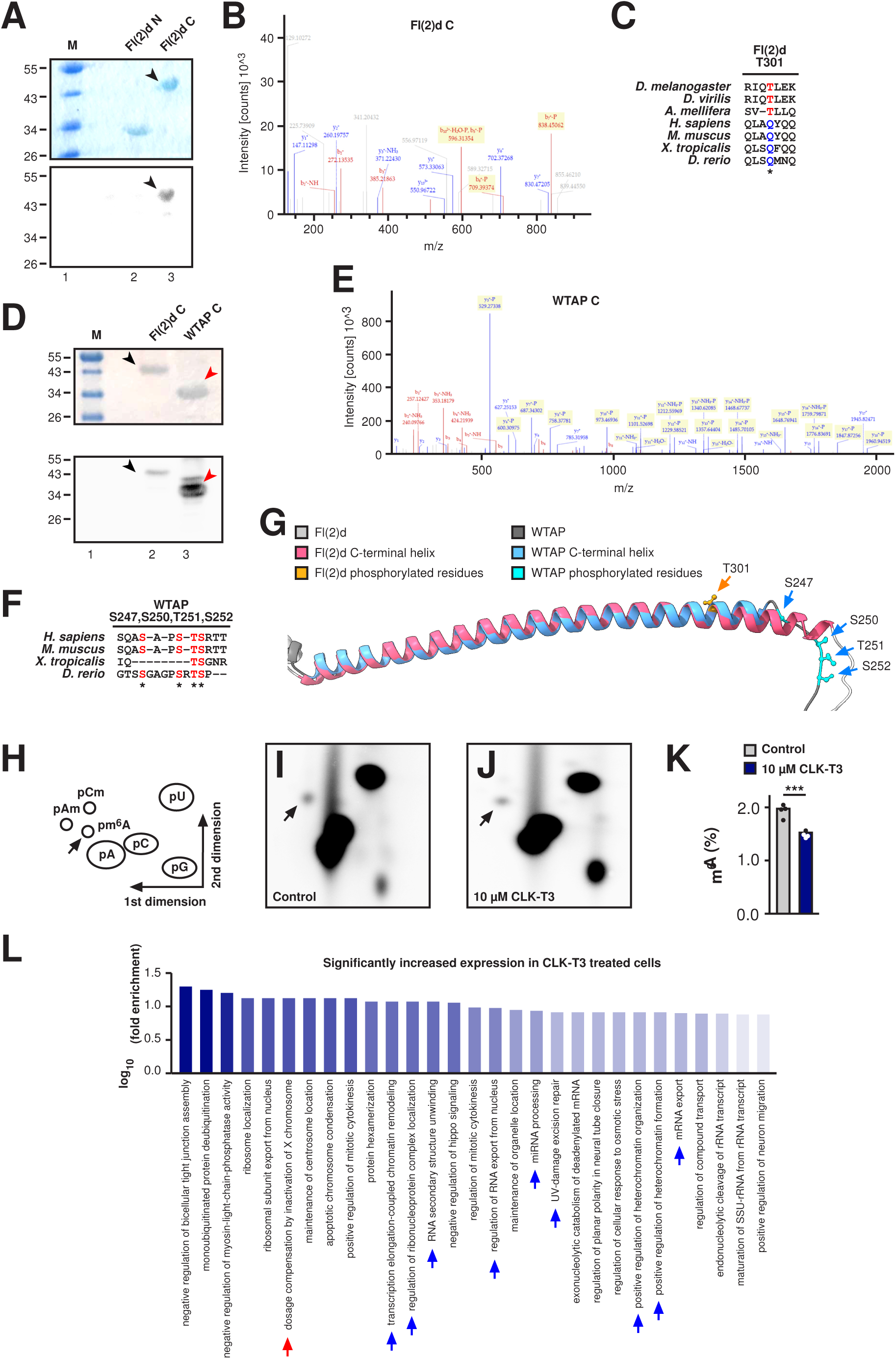
DOA/CLK2 phosphorylation of Fl(2)d and WTAP is required for m^6^A methylation in *Drosophila* and humans. A) Radio-labelling of Fl(2)d-N and -C termini through CLK2 phosphorylation run on a denaturing gel. Coomassie-stained proteins are shown on the top and ^32^P-phosphorylated bands on the bottom gel. B and C) LC-MS/MS analysis of Fl(2)d-C phosphorylation at residue T301 (B) and alignment of the phosphorylated Fl(2)d T301 between insects. D) Radio-labelling of Fl(2)d-C and the WTAP C-terminal helix through CLK2 phosphorylation run on a denaturing gel. Coomassie-stained proteins are shown on the top and ^32^P-phosphorylated bands on the bottom gel. E and F) LC-MS/MS analysis of WTAP-C phosphorylation at residues S247, S250, T251, and S252 (E) and alignment of the phosphorylated WTAP residues between vertebrates. G) Structural alignment of the Fl(2)d/WTAP C-terminal helix with the phosphorylated Fl(2)d T301 residue and WTAP S247, S250, T251, and S252 residues indicated. H-K) Schematic representation of a TLC depicting non-methylated and methylated nucleotide positions (H), and TLCs displaying m^6^A in untreated (I) and 10 µM CLK-T3 24 hour treated Hek293T cells (J). Quantification of m^6^A compared to A is shown in (K). Statistical significance was calculated from unpaired student t-tests, indicated by asterisks (*** p≤0.001). L) Gene ontology analysis from 3’ RNA-seq in HeLa cells treated with 1 µM CLK-T3 for 8 hours (Faraway et al. 2023). Significantly increased expression changes enriched in m^6^A- regulated pathways are indicated with arrows.

We next tested whether CLK2 phosphorylates WTAP, the human homologue of Fl(2)d. Mass spectrometry analysis of post-translational modifications on Fl(2)d C-terminal fragment revealed phosphorylation of four residues, serine 247 (S247), S250, T251, and S252, which are conserved in vertebrates (Fig. 4B and C). The prominent alpha-helix in WTAP is conserved in *Drosophila* Fl(2)d evident from Alphafold structural modelling. However, the alpha-helix adjacent regions likely comprising an interaction interface have diverged. Despite the fundamental function of the core MACOM complex members WTAP/Fl(2)d, Virma/VIR and Flacc their evolutionary conservation is low (Balacco and Soller 2019; Ensinck et al. 2023). Taken together, the CLK2 phosphorylation sites have shifted to the end of the alpha-helix, while the phosphorylation site from insects became fixed into a conserved glutamine, which is a partial phosphomimetic for phosphorylated threonine (Fig. 4C-G, Supplementary Fig. 6).

### The DOA human homologue CLK2 is required for m^6^A methylation in humans

Since DOA is highly conserved in humans and other organisms (Fig. 1A, Supplementary Fig. 1), we wanted to test whether its homologue CLK2 is required for m^6^A writer complex activity in human cells. Therefore, we incubated HEK293T cells with the CLK1-3 inhibitor CLK-T3 for 24 h and determined m^6^A levels in poly(A) RNA by TLCs (Funnell et al. 2017; Faraway et al. 2023). We observed a 25% reduction of m^6^A in the CLK- T3 treated cells compared to untreated cells (Fig. 4H-K).

Although sex determination between flies and mammals is different, m^6^A plays key roles in this pathway across species (Roignant and Soller 2017). Since m^6^A in non-coding RNA XIST is required for dosage compensation in humans we determined whether DOA has a role in this pathway (Patil et al. 2016). Accordingly, we analysed 3’ end sequencing data for cells treated with the CLK2 inhibitor CLK-T3 (1 µM for 8 hours) (Faraway et al. 2023). Gene ontology analysis for all genes with significant changes in expression revealed a significant enrichment of genes involved in dosage compensation, as well as other known m^6^A-regulated pathways (Fig. 4L).

Taken together, these results suggest a conserved role for DOA/CLK2 in regulating m^6^A writer complex activity through phosphorylation of WTAP/Fl(2)d identified through its key role in *Drosophila* Sxl auto-regulation required for sex determination and dosage compensation (Supplementary Fig. 7).

## Discussion

In this study we identified a novel role for DOA in regulating female-specific *Sxl* alternative splicing essential for female sexual differentiation and X chromosome dosage compensation. DOA is required for m^6^A mRNA methylation and reinforces autoregulation of *Sxl* alternative splicing by increasing the activity of the m^6^A mRNA methylation writer complex through phosphorylation of Fl(2)d. This view is supported by genetic dosage alteration experiments of writer components through a read-out in dosage compensation leading to female lethality, or after removal of the Sxl target *msl-2* in sexual transformations. Moreover, knockdown of CLK2 in human cells reduces m^6^A levels and affects many m^6^A regulated processes including genes involved in human dosage compensation, indicating a conserved regulatory mechanism across species with implications for developmental processes and disease.

The m^6^A methyltransferase complex consists of two conserved sub-complexes MAC and MACOM that need to interact for m^6^A mRNA methylation in animals, plants and fungi (Knuckles et al. 2018) (Balacco and Soller 2019; Brodersen and Arribas-Hernández 2024). MAC comprises the catalytic core of the m^6^A complex, consisting of the Mettl3-Mettl14 heterodimer with little methylation activity on its own in vitro (Su et al. 2022). For full methylation activity, the auxiliary MACOM complex is required as well. A mechanistic understanding of its requirement, however, remains to be established. The large size of the m^6^A writer complex implicates that m^6^A deposition is regulated process through cellular signalling, but how post-translational modifications affect complex activity is unclear.

We genetically determined that the molecular target of the highly conserved kinase DOA is Fl(2)d through its key role in regulating Sxl autoregulation required for *Drosophila* sex determination and dosage compensation. Here, DOA phosphorylates Fl(2)d in a highly conserved structured alpha-helix at a single threonine residue.

DOA has previously been attributed a role in the regulating SR protein phosphorylation to alter sexual differentiation through TRA, TRA2, and RBP1 in *Drosophila* (Du et al. 1998). Hence, DOA can operate at multiple levels in the sex determination pathway to coordinate sexual differentiation. Although the human homologue of DOA, CLK2 phosphorylates SR proteins (Colwill et al. 1996), whether this activity is coordinated with the m^6^A pathway remains to be determined.

While in *Drosophila* phosphorylation of Fl(2)d by DOA is required for m^6^A mRNA methylation by the writer complex, this seems to be less stringent in human HEK293T cells as pharmacological inhibition of CLK2 did not result in complete loss of m^6^A. To fully understand the extend of this regulation, however, a vertebrate animal model is required.

A more elaborated regulation of m^6^A writer complex activity in humans, however, is suggested by phosphorylation of three amino acids in WTAP compared to only one in *Drosophila* Fl(2)d. In vertebrates, the highly conserved phosphorylation site in insects has changed to an evolutionary fixed glutamine, which is phosphomimetic-like for phosphorylated threonine. In addition, CLK2 phosphorylation is shifted further away from the core MACOM. Although the full structure of the MACOM is not known (Su et al. 2022), the phosphorylation sites likely are in an interaction interface with another MACOM member. Delineating mechanistic insights into the exact role of phosphorylation from the partial complex structure obtained by Cryo-EM is complicated by the low evolutionary conservation of the three MACOM members Fl(2)d/WTAP, Vir/VIRMA, and Flacc. These three MACOM members are characterised by extended low complexity regions requiring further structural and biophysical information to understand the impact of phosphorylation. Potentially, polymorphisms associated with human disease could aid a mechanistic understanding.

The exon junction complex (EJC) has been shown to interfere with m^6^A deposition around exon junctions in coding sequences by association with the transcription and export (TREX) complex in nuclear mRNPs for export from the nucleus (Yang et al. 2022; He et al. 2023; Luo et al. 2023; Uzonyi et al. 2023; He and He 2024). In the absence of EJC components, m^6^A becomes deposited in some short exons and decreases mRNA stability (Yang et al. 2022; He et al. 2023; Luo et al. 2023; Uzonyi et al. 2023; He and He 2024). Although m^6^A is not installed in all possible places in the absence of the EJC, and is not enriched in 5’ UTRs where the EJC is not present, a model has been proposed that m^6^A deposition is passive (Yang et al. 2022; He et al. 2023; Luo et al. 2023; Uzonyi et al. 2023; He and He 2024). Accordingly, the EJC competes with binding of the m^6^A writer complex to prevent m^6^A installation (Yang et al. 2022; He et al. 2023; Luo et al. 2023; He and He 2024). However, the m^6^A writer complex is recruited co-transcriptionally and in *Drosophila Sxl* m^6^A is deposited before splicing (Haussmann et al. 2016), which has also been described for many human genes (Louloupi et al. 2018; Tang et al. 2024). Although the EJC is deposited 24 nucleotides upstream of splice junctions after splicing its role in competing with the m^6^A writer complex likely originates much earlier. In fact, the EJC is recruited by the splicing factor CWC2 (Alexandrov et al. 2012) co-transcriptionally suggesting a role in exon definition which occurs shortly after short exons are transcribed (Keren et al. 2010; Ule and Blencowe 2019). Accordingly, splicing defects have been identified in EJC mutants that are linked to exon definition (Ashton-Beaucage et al. 2010; Roignant and Treisman 2010; Akhtar et al. 2019).

An active role for m^6^A deposition is further indicated by cell-type-specific regulation of m^6^A-methylated mRNAs (Flamand and Meyer 2022; Perlegos et al. 2024). For instance, mRNA localisation to neuronal projections can be attributed to specific m^6^A sites and their mutation abolishes localization (Flamand and Meyer 2022). Likewise, our findings also support an active role of m^6^A methylation. We observed m^6^A-induced cell- and tissue-specific sexual transformations.

Competition between RNA binding proteins for binding sites constitutes one of the key principles in the regulation of alternative mRNA processing (Soller 2006). However, this thus not mean that individual processes such as m^6^A methylation are passive. Our data clearly show that m^6^A deposition is regulated by cellular signalling. The diverse phenotypes from genetic interaction experiments in *Drosophila* sex determination together with the numerous post-translational modifications found in m^6^A writer proteins further point towards complex regulation by cellular signalling (England et al. 2022).

## Methods

### *Drosophila* stocks, genetics, immunostainings and imaging

*D. melanogaster* CantonS and *w^1118^* were used as the wild type control. *Doa^HD^*, *Doa^DEM^* and *Doa^EMS-2^*were kindly provided by Leonard Rabinow. *Doa^MB^* (BDSC 23390), Df(3R)BSC789 (BDSC 27361), *UASDoa* (BDSC 20286), and *Doa^Dp^* (89894) were from Bloomington Stock Centre. All other alleles used in this study have been described previously (Haussmann et al. 2016; Knuckles et al. 2018; Bawankar et al. 2021). For clonal analysis, *Doa^DEM^* and *Doa^EMS-2^* were recombined onto FRT82B and clones were generated with hsflp attP40; FRT82B ubiGFP by heat-shock in early larvae.

Drosophila cultures were maintained at 25°C in plastic vials containing standard cornmeal/yeast-rich medium (1% agar, 2% yeast, 7% dextrose, 8% cornmeal w/v and 2% Nipagin from a 10% solution in ethanol) with a 12:12 hour light-dark cycle. Percentage female viability was calculated by dividing the total number of females by the total number of males. Significance was calculated through chi-squared statistical tests with FDR corrected significance value of p<0.05. Fly images were taken using a ZEISS AxioCam ICc 1 with Stemi 2000-CS microscope with the ZEN 2012 software.

For analysis of synapses at NMJs of third instar wandering larvae we selected wild type, *Mettl3^null^* homozygous, and *Doa^EMS-2^/Doa^DEM^* females. Larvae were dissected in PBS and fixed in Bouin’s solution (Sigma-Aldrich, HT10132) for 5 min and washed in PBT (PBS with 0.2% BSA and 0.1% TritonTM X-100 (Sigma, T8787)). Samples were incubated with primary antibodies rabbit anti-HRP (1:250, 323 005 021, Jackson ImmunoResearch) and mouse anti- NC82 (1:100, DSHB) at 4 °C overnight. Following washing in PBT tissues were incubated overnight at 4 °C with secondary antibodies conjugated with Alexa Fluor 488 (anti-rabbit) or Alexa Fluor 546 (anti-mouse). NMJs were mounted in Vectashield (Vector Labs) and scanned with Zeiss Imager.M2 ApoTome.2. Images were processed using FIJI.

### RNA isolation, mRNA purification and RT-PCR

Total RNA was extracted from using Tri-reagent (SIGMA), with subsequent removal of DNA through treatment with DNase I (New England Biolabs). Fly heads and thoraxes, and abdomens were dissected from 3- to 5-day old flies and homogenised in Tri-reagent followed by RNA isolation. Poly(A) mRNA was isolated via two rounds of purification with Dynabeads Oligo d(T)25 according to the manufacturer (New England Biolabs) from 5 flies or 10 head/thoraces.

Preparation of cDNA via reverse transcription was performed using Superscript II (Invitrogen) using an oligo-dT17V primer as previously described (Haussmann et al. 2016). PCR for alternative *Sxl* exon 3 was performed for 40 cycles with 1 μl of cDNA *Sxl* F2 (ATGTACGGCAACAATAATCCGGGTAG) and *Sxl*R2 (CATTGTAACCACGACGCGACGATG). Experiments included at least two biological replicates.

### TLC analysis of m^6^A levels

TLC analysis was performed on half of the eluate from polyA selection described above (about 50 ng). Simultaneous digestion of poly(A) mRNA with RNase T1 (Fermantas) and 5’end-labelling with 0.5 μl [γ-32P] ATP (6000 Ci/mmol, 25 μM, Perkin-Elmer) and 10 U of T4 PNK (NEB)was performed in T4 PNK buffer for 2 hours at 37 °C. Following ethanol precipitation (2.5 Volumes) in the presence of 0.3 M NaAc and washing twice with 70% ethanol, RNA was resuspended in 10 μl of 50 mM sodium acetate (pH 5.5) and digested with nuclease P1 (SIGMA) for 2 hours at 37 °C. Then 1 μl of the digest RNA sample was spotted on cellulose F TLC plates (20 × 20 cm, Merck) and run for the first dimension in a solvent system of iso-butyric acid:NH4OH(28-32%, 7.7M)/water (62.56:2.44:35, v/v/v), and for the second in isopropanol:HCl:water (70:15:15, v/v/v). Experiments were performed from at least two biological replicates and nucleotide identities were assigned as previously described (Haussmann et al. 2016). Storage phosphor screens (K-Screen, Kodak) and Molecular Imager FX were used to quantify spot intensities with QuantityOne (BioRad).

### Cell culture and CLK-T3 inhibitor treatment

Human embryonic kidney (HEK)-293T cells were cultured in Dulbecco’s Modified Eagle Medium (DMEM, Merck) supplemented with 4 mM L-glutamine, 10% fetal bovine serum (FBS, Thermo Fisher Scientific), 100 U/ml penicillin, and 100 μg/mL streptomycin (Merck). The inhibitor CLK-T3 (Sigma, SML2649) was dissolved to 5 mM in DMSO. Cells were treated with 10 μM CLK-T3 and control cells were treated with equivalent volume of DMSO for 24 hours.

### Cloning, recombinant protein expression, and purification

Plasmids used for Fl(2)d and WTAP recombinant protein expression were constructed by cloning the corresponding cDNA into pGex vector containing an N-terminal GST-tag and recombinant proteins were made as described (McQuarrie and Soller 2024). The N-terminal fragments for Fl(2)d and WTAP were amplified with primers Fl(2)d NF1 (gtGATTACAAAGACGACGATGACAAgcttGCTCAGCAATGCGCGGACGCCCAGCGA C) and NR1 (cagtcacgatgaattgcggccgctcTagattaAGCGTCCTCGTGAATGGTCTCTAG), and the C-terminal fragment with primers Fl(2)d CF1 (gtGATTACAAAGACGACGATGACAAgcttCAGCAAGAACTAAAGACCACACGCGAT C) and CR1 (cagtcacgatgaattgcggccgctcTagattaGGTGGAGTAGTCGACTGCTCCGCTG), and WTAP NF1 (agtGATTACAAAGACGACGATGACAAgcttGGACGTATTGCACAACTTGAAGCAGA G) and NR1 (tcagtcacgatgaattgcggccgctcTagattaACCCTGTACATTTACACTTGAGTC), respectively. Recombinant protein was cleaved off the GST-moiety with PrecissionProtease (Amersham) and stored in protease cleavage buffer (50 mM Tris, 150 mM NaCl, 0.5 mM DTT, 0.5 mM EDTA).

### Phosphorylation assays and LC-MS/MS analysis

Phosphorylation assays were performed using the CLK2 Kinase Enzyme System kit (VA7414, Promega) with recombinant proteins for Fl(2)d and WTAP. For each protein, 10 μg was incubated with 0.5 μl CLK2 (0.05 μg, C58-11G-10, Promega), 0.5 μl [γ-32P] ATP (6000 Ci/mmol, 25 μM, Perkin-Elmer), and 1 mM DTT in 1x Reaction Buffer A (K03-09, Promega) to a total volume of 25 μl for 1 hour at room temperature. Protein bands were separated on SDS denaturing gels and imaged using storage phosphor screens (K-Screen, Kodak) and Molecular Imager FX. Phosphorylated bands were analysed using Quantity One 1-D (Bio-Rad) as previously described (Ustaoglu et al. 2024).

Samples were prepared for LC-MS/MS as above in the presence of 1 mM ATP without denaturing gel separation. Peptide concentration and separation was performed using an UltiMate 3000 HPLC series (Dionex, Sunnyvale, CA USA). Samples were trapped on an Acclaim PepMap 100 C18 uPrecolumn (5 µm, 100Å, 300 µm i.d. x 5 mm, Dionex) following separation with a Nano Series C18 PepMap100 column (3 µm, 100Å, 75 µm i.d. x 15 cm, Dionex). The gradient ranged from 3.2% to 44% solvent B (0.1% formic acid in acetonitrile) over 30 minutes. Peptides were eluted (∼350 nL/min) using a Triversa Nanomate nanospray source (Advion) into a QExactive HF (Thermo Fisher) mass spectrometer, controlled by Xcalibur 4.0. Data-dependent acquisition alternated between full FT-MS scans (m/z 360–1600) and high-energy collision dissociation (HCD) MS/MS of the top 20 ions. Survey scans had a resolution of 120,000 at m/z 200 with AGC set to 3x106. HCD MS/MS scans had a resolution of 15,000, a normalized collision energy of 28, and an AGC target of 1x105, using a 1.2 m/z isolation window for multiply charged precursors. Spectra were collected over 56 minutes with a 20-second dynamic exclusion. The UniProt database was used to identify samples using Proteome Discoverer 2.2. Variable modifications include deamidation (N, Q), oxidation (M), and phosphorylation (S, T, Y), with a 10-ppm precursor mass tolerance and a 0.02 Da MS/MS tolerance.

## Supplementary information

Supplementary information includes Supplementary Dataset 1.

## Supporting information

Supplementary Results

## Acknowledgements

We thank the Bloomington Stock Centre and Lenny Rabinow for fly lines, Mike Tomlinson and Connie Koo for help with cell culture, Frannie H. S. Stephens for reagents, Irmgard U. Haussmann and Alper Akay for comments on the manuscript. This work was supported by the Medical Research Council (MR/N013913/1) to D.W.J.M. and the Biotechnology and Biological Science Research Council and the Leverhulme Trust to M.S.

## Author contributions

D.W.J.M. conceived the project. D.W.J.M. and M.S. directed the project. D.W.J.M. performed genetic experiments, cell culture, immunostaining, biochemical experiments, evolutionary analyses, differential gene expression, and structural modelling. W.B. and D.W.J.M. performed cloning and protein purification experiments. D.W.J.M. and M.S wrote the manuscript. All authors read and approved the final manuscript.

## Availability of data and materials

All data generated or analysed during this study are included in the supplementary information files.

## Declarations

### Ethics approval and consent to participate

Not applicable.

### Consent for publication

Not applicable.

