## Supplementary Results for "DOA/CLK2 phosphorylates Fl(2)d/WTAP to enhance m^6^A mRNA methyltransferase complex activity"

To examine the role of DOA in regulating the m$^6$A methyltransferase complex, we tested female viability in a range of sensitised backgrounds with m$^6$A complex components removed in either heterozygous or homozygous conditions. Here we observed that female viability of the sensitised background *vir$^{2F}$*/ *vir$^{ts}$* was ~30-40%, and that by removing one copy of *Hakai*, *nito*, or *Doa* female viability remained low, but that females were still viable (Supplementary Fig. 4). Removal of one copy of both *Hakai* and *nito* in the *vir$^{2F}$*/ *vir$^{ts}$* background resulted in female lethality, likely resulting from impaired m$^6$A methyltransferase stoichiometry (Supplementary Fig. 4). Using the homozygous viable *Doa$^{MB}$* mutant (~95% female viability), we tested female viability in the sensitised *vir$^{2F}$*/ *vir$^{ts}$* background and found that females were no longer viable, mimicking the observed phenotype for removal of one copy of both *Hakai* and *nito* in the same background (Supplementary Fig. 4). Further, by removing one copy of *Doa$^{MB}$* and *Hakai* we observed that these females were no longer viable, again mimicking the phenotype associated with m$^6$A complex sub-stoichiometry. These results support that DOA has a role in regulating m$^6$A methyltransferase complex activity.

**Supplementary Fig. 1: Amino acid alignment of DOA/CLK2.**

Amino acid alignment of DOA and CLK2 between *D. melanogaster* and humans, respectively. Identical amino acids are highlighted (red), complementary amino acid substitutions based on the BLOSUM-62 substitution matrix (orange), and non-complementary amino acid substitutions (unmarked) are indicated.

**Supplementary Fig. 2: DOA regulates cell-specific female differentiation in the abdomen.**

A) Quantification of male-specific pigmentation and sexual development in females in sensitised *Sxl/+*; $da^{Df}$/+ backgrounds with heterozygous *Doa* alleles.

B-D) Representative images of wild-type male and female pigmentation (B and C) and *Sxl/+*; $da^{Df}$/+; $Doa^{DEM}$/+ male-specific pigmentation in females (D). The scale bar (B) is 200 μm.

**Supplementary Fig. 3: DOA and Mettl3 regulate synaptic bouton numbers in motor neurons.**

A-B) Representative images (A) and quantification (B) of female third instar larval NMJs from muscle 13 synapses for wild-type, $Mettl3^{null}$, and $Doa^{EMS\text{-}2}/Doa^{DEM}$ genotypes. Motor neurons were stained with anti-HRP (magenta) and synaptic boutons with anti-NC82 (green, the merged colour is white). Statistically significant differences from the wild-type control were calculated from unpaired student t-tests following Bonferroni correction are indicated by asterisks (**** $p \leq 0.0001$). The scale bar (A) is 10 μm.

**Supplementary Fig. 4: Female viabilities of $m^6A$ writer and DOA mutants in combination.**

Quantification of female viability in different combinatorial allelic backgrounds. Statistically significant differences from the controls were calculated from chi-squared tests following FDR correction are indicated by asterisks (*** $p \leq 0.001$, **** $p \leq 0.0001$).

**Supplementary Fig. 5: Analysis of m$^6$A complex stability in *Doa* null cells.**

A) Clonal analysis of Fl(2)d protein levels in *Doa* null and wild type cells. DAPI is shown in blue (A), ubiquitin GFP in green and *Doa*$^{EMS\text{-}2}$ homozygous cells in black (A'), and Fl(2)d in red (A''). No discernible reduction of Fl(2)d was observed.

**Supplementary Fig. 6: Fl(2)d/WTAP phosphorylated positions in the partial 'warhorse' m$^6$A writer complex structure.**

The structure depicts the partial 'warhorse' m$^6$A writer complex, highlighting the phosphorylated residues of Fl(2)d/WTAP identified by mass spectrometry. Phosphorylation sites are marked in orange (Fl(2)d T301) or yellow (WTAP S247), mapped onto the structural model to indicate their spatial distribution within the complex.

**Supplementary Fig. 7: Model showing the function of the m$^6$A complex in the presence and absence of DOA.**

A model of DOA-regulate m$^6$A methylation in *Drosophila*.

### Supplementary Figure 1

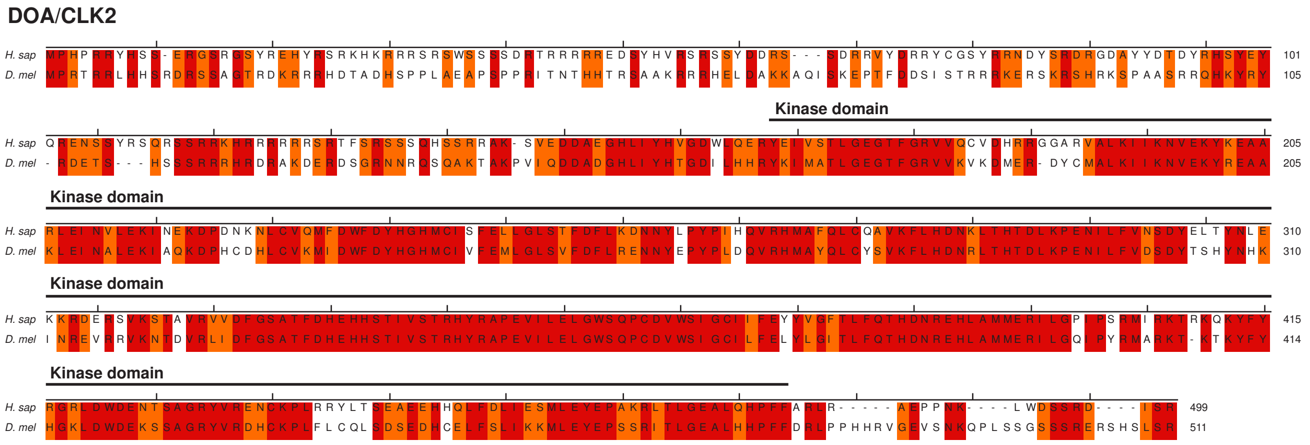

### Supplementary Figure 2

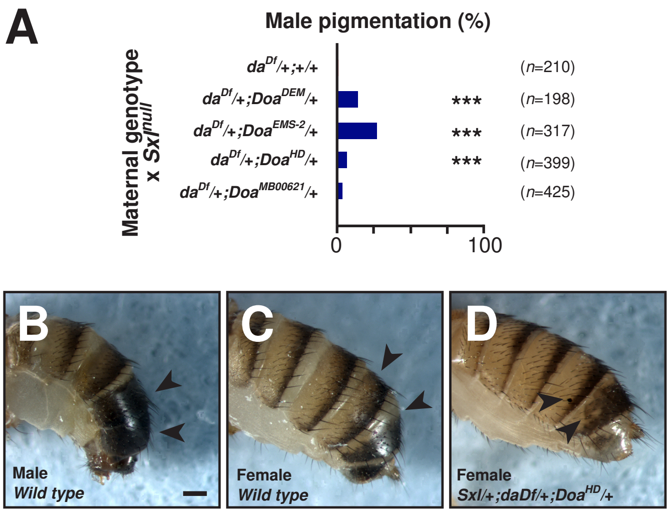

### Supplementary Figure 3

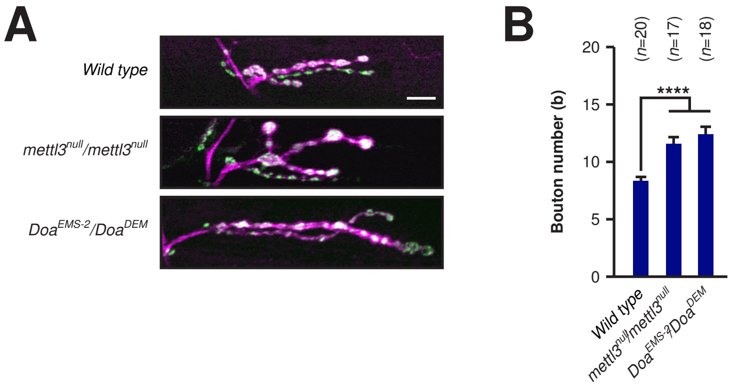

### Supplementary Figure 4

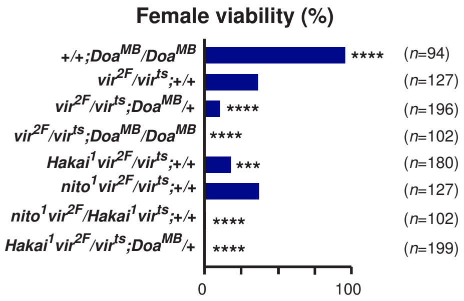

### Supplementary Figure 5

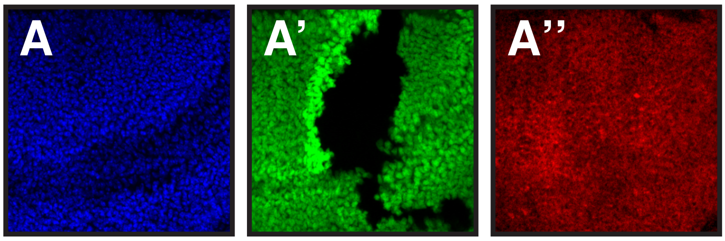

### Supplementary Figure 6

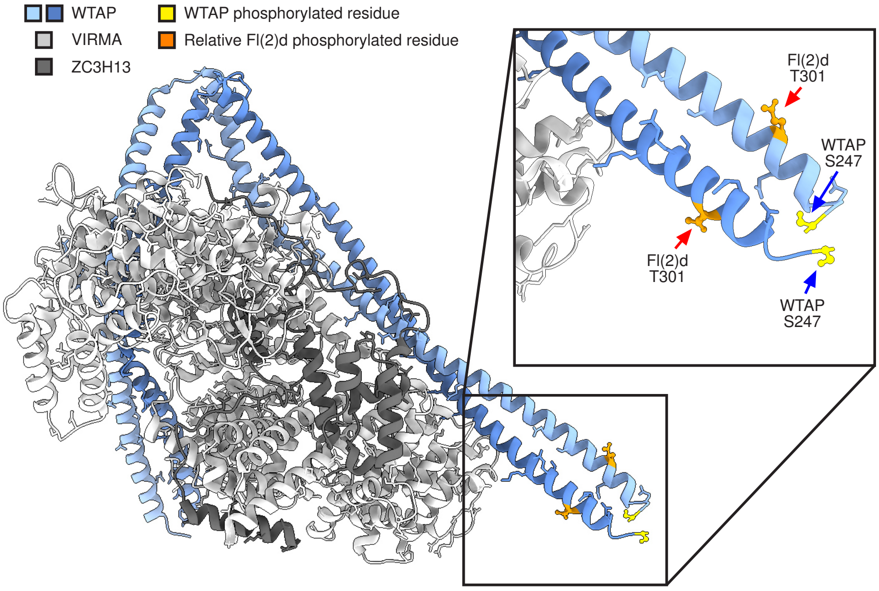

### Supplementary Figure 7

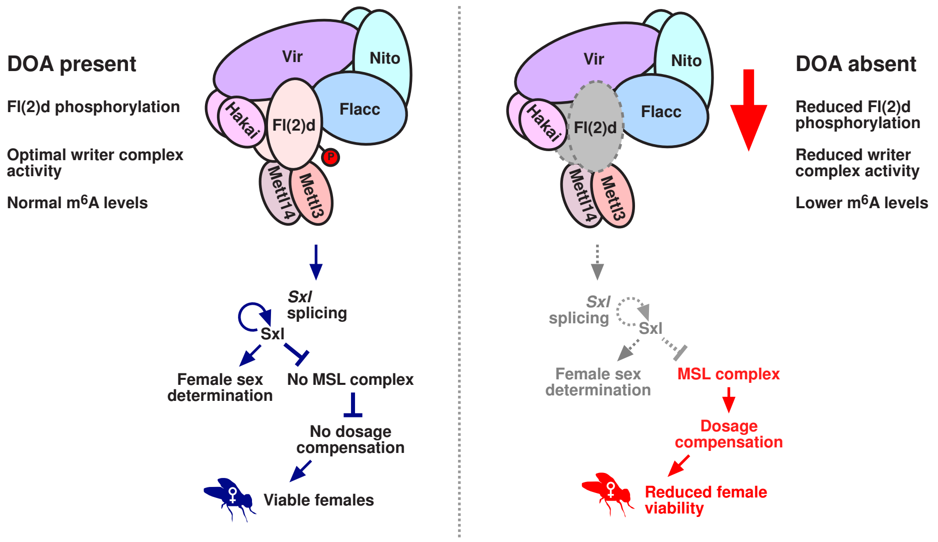
